# Endothelial Cell Specific Thioredoxin 1-Dependent Signaling in Oxygen-Induced Retinopathy

**DOI:** 10.64898/2026.11.04.763768

**Authors:** Henry H. Song, Wenjing Wu, Daniyal Khan, Hua Zhong, Mathew S. Meaders, Paul T. Pierce, Peter F. Vitiello, Lynette Rogers, Trent E. Tipple, Faizah N. Bhatti

**Author notes:** Corresponding Author: Faizah N. Bhatti, MD, MS, Departments of Pediatrics, Ophthalmology & Cell Biology, The University of Oklahoma Health Campus, 800 Research Parkway, Room 464, Fourth Floor, Oklahoma City, OK 73104.

## Abstract

**Purpose:** Retinopathy of prematurity (ROP) is a leading cause of childhood blindness, characterized by oxidative stress and dysregulated angiogenesis in preterm infants with immature antioxidant systems. This study aimed to investigate the specific role of endothelial cell-specific thioredoxin 1 (TXN1), a crucial oxidoreductase, in oxygen-induced retinopathy, hypothesizing that the absence of endothelial TXN1 directly impairs angiogenesis and exacerbates hyperoxia-induced injury.

**Methods:** We generated tamoxifen-inducible, endothelial cell-specific *Txn1* knockout mice (*Txn1-iECKO*) and utilized the oxygen-induced retinopathy (OIR) model, exposing litters to 85% oxygen from postnatal day 7 (P7) to P12. Retinal vascular morphology during vaso-obliteration and neovascularization phases, endothelial cell proliferation, and apoptotic cell death were assessed. Furthermore, human retinal microvascular endothelial cells (HRMECs) with TXN1 knockdown were subjected to hyperoxic conditions, followed by comprehensive gene expression profiling using RT^2^ Profiler PCR arrays and Ingenuity Pathway Analysis.

**Results:** Deletion of endothelial *Txn1* significantly enhanced vascular regression during the hyperoxic vaso-obliteration phase at P12 and markedly exacerbated pathological neovascularization at P17. *Txn1*-deficient retinas exhibited significantly impaired endothelial cell proliferation during the vaso-obliteration phase, while apoptotic cell death remained unchanged. Transcriptomic analysis of TXN1-deficient HRMECs under hyperoxia revealed an enhanced activation of NRF2-mediated oxidative stress response pathways.

**Conclusions:** These findings identify endothelial TXN1 as an essential and oxygen-dependent regulator of vascular development in ischemic retinopathy, primarily influencing endothelial proliferation and redox signaling. This suggests TXN1’s crucial protective role and highlights its therapeutic potential to mitigate ROP progression by modulating endothelial cell responses to oxidative stress.

## INTRODUCTION

Preterm birth rates have been increasing over the last two decades, posing a significant health burden not only for the US but globally. Resuscitation at earlier gestational ages is associated with increasingly complex disruption and dysplasia at the cellular, organ, and systemic levels. Retinopathy of prematurity (ROP) is one such complication and the leading cause of childhood blindness.^1,2^ The risk of ROP is inversely related to birth weight and gestational age. It is influenced by nutrition and immaturity of immune and redox pathways, which serve as the first line of defense against environmental challenges.^3,4^ During ROP, retinal vascularization occurs in two distinct phases: hyperoxia-induced arrest of normal retinal vascular development, followed by hypoxia-driven pathological neovascularization. This complex, multifactorial process involves dysregulated cytokine expression and aberrant signaling pathways.^1^

Atmospheric oxygen is hyperoxic relative to the *in-utero* environment. The abrupt postnatal transition to hyperoxia elicits a potent oxidative stress response, characterized by overproduction of reactive oxygen species and inhibition of crucial hypoxia-dependent pathways. The fetus prepares for this shift through enhanced antioxidant systems like thioredoxin,^5,6^ glutathione,^7^ and superoxide dismutase,^8^ which are critical for maintaining cellular redox balance by scavenging ROS.^9^ However, preterm infants are born with underdeveloped retinal vasculature and immature antioxidant systems, making them vulnerable to postnatal hyperoxia.^10,11^ Supplemental oxygen therapy, often administered for respiratory distress, imposes an additional hyperoxic insult, correlating with increased ROP risk and severity. This inherent antioxidant deficiency exacerbates oxidative stress and cellular damage, intensifying ROP severity by compromising the cellular redox state. ^7,12,13^

Thioredoxin 1 is a ubiquitously expressed oxidoreductase vital for homeostatic maintenance and cellular response to oxidative stress. TXN1 regulates cellular redox by reducing oxidized proteins, contributing to redox signaling, and directly quenching hydrogen peroxide.^5^ Under normoxic conditions, apoptosis signal-regulating kinase 1 (ASK1) is sequestered by reduced TXN1. In response to oxidative stress, thioredoxin-interacting protein binds directly to TXN1^14^ and disrupts ASK1-TXN1 inhibitory binding, triggering ASK1 activation and apoptosis.^15,16^ TXN1 confers protection against ROP in murine models of perinatal hypoxia-ischemia^17^ and oxygen-induced retinopathy^18^, and plays a regulatory role in proinflammatory and angiogenic signaling. The role of TXN1 in ROP progression has not been investigated in a cell type-specific *in vivo* context. We hypothesize that TXN1 deficiency in endothelial cells directly impairs angiogenesis, exacerbated by hyperoxia. This study aims to determine if retinal EC-expressed TXN1 protects against oxygen-induced pathological vascularization, and to elucidate TXN1’s protective mechanisms against oxidative stress-induced retinal damage. Specifically, we focus on its role in modulating key signaling pathways in pathological angiogenesis and oxidative stress, thereby providing a deeper understanding of its therapeutic potential in mitigating ROP progression.

## METHODS

### Animals

All animal procedures were approved and performed under the guidelines and regulations of the OU Health Campus Institutional Animal Care and Use Committee (protocol# 24-049-H) and adhered to standards set by the Association for Research in Vision and Ophthalmology (ARVO). The OU Health Campus facility is fully accredited by AALAC, International, has an Animal Welfare Assurance on file with the Office of Laboratory Animal Welfare (A3165-01), and is a licensed research facility under the authority of the United States Department of Agriculture (73-R-0007).

### Generation of mice

*Txn1* floxed mice were generated using CRISPR/Cas9-mediated homologous recombination. Guide RNAs (gRNAs) were designed to insert *loxP* sequences into *Txn1* at intron 1 (5’-ACTCACGCTGGAGGCCAACA<u>TGG</u>-3’) and downstream of the 3’ UTR (5’-GAAATACAATGTCCCACTCC<u>AGG</u>-3’), with the PAM sequence underlined. Genomic DNA was prepared using AccuStart II Mouse Genotyping Kit (Quanta Biosciences), and sequence analysis and genotyping were performed using the primer sets for 5’ *loxP* site on intron 1 and 3’ *loxP* site on the 3’ UTR of the *Txn1* gene. Primers 5’-CCACTTTATGTGGTTGCTTTGGA-3’ and 5’-CTCCTCGCACCCATTAATTTCAA-3’ were used for the 5’ region, yielding amplicons of 193 bp for the wild-type allele and 227 bp for the floxed allele. For the 3’ region, primers 5’-GGCACAAGTTCTCCATAAGCCTGGCA-3’ and 5’-TCTGGGGCCAAAGGTTCAAATCCCA-3’ were used, producing amplicons of 204 bp for the wild-type allele and 238 bp for the floxed allele. A tamoxifen-inducible *Rosa26-CreER* mouse line (Jackson Laboratory) was used to verify Cre-mediated deletion of *Txn1* floxed alleles after tamoxifen treatment. TXN1 protein and gene expression were assessed by immunoblotting and RT-qPCR, respectively, in lung and brain tissues (TXN1 antibody; Cat. #2429; Cell Signaling Technology). *Cdh5-CreERT2* mice^19^ were kindly provided by Dr. Ralf H Adams at Max Planck Institute for Molecular Biomedicine (Münster, Germany). To generate inducible, EC-specific *Txn1* knockout mice, *Cdh5-CreERT2* mice were crossed with *Txn1^fl/fl^* mice to generate *Txn1^fl/+^;Cdh5-CreERT2^+^* mice. These mice were bred with *Txn1^fl/fl^* mice, yielding *Txn1^fl/fl^;Cdh5-CreERT2^+^* mice, which served as EC-specific *Txn1* knockout mice following tamoxifen administration. Tamoxifen dosage, administration method, and endothelial-specific Cre recombinase activity, particularly in the retina, were validated using *mT/mG;Cdh5-CreERT2* reporter mice.

### Mouse Oxygen-induced retinopathy (OIR) Model

The OIR model is well-established for recapitulating key pathological features of ROP, including vascular arrest and subsequent neovascularization.^20,21^ To induce Cre recombinase activity in the *Cdh5-CreERT2* mice, tamoxifen (Sigma, T5648) (in peanut oil at 25 mg/mL) was administered by oral gavage at 50 µg on postnatal days 5 (P5), P6, and P7. Litters, along with their dams, were exposed to either 75% oxygen or 21% room air (control) for 5 days, from P7 to P12 (hyperoxia phase), after which they were returned to room air until P17 (hypoxia phase). Eyes were enucleated, and whole retinas were isolated by microdissection at P7, P12, or P17, for subsequent analysis.

### Immunohistochemistry (IHC)

Retinal vascular morphology was examined by fluorescent IHC staining as previously described.^21^ Enucleated eyes were fixed in 4% PFA for one hour at room temperature, followed by dissection and isolation of the intact retina. Retinal tissues were blocked overnight at 4°C with PBS containing 5% BSA, 5% normal horse serum, 0.5% Triton X-100, and 0.02% NaN_3_. For blood vessel staining, the primary antibody used was Alexa Fluor 488-conjugated isolectin GS-IB4 (Invitrogen, Cat. #I21411) or hamster anti-mouse CD31 antibody (Millipore, Cat. #MAB1398Z). After incubating with the primary antibody, samples were washed three times for 15 minutes each in PBS containing 1% BSA, 1% normal horse serum, and 0.1% Triton X-100 at room temperature. For the CD31 primary antibody, the secondary antibody used was Alexa Fluor 594-conjugated goat anti-Armenian hamster IgG (Jackson ImmunoResearch, Cat. #127-585-099). Retinal tissue was then cut into a clover leaf motif by microdissection and flat-mounted onto glass slides. Images were captured using either a Zeiss Axioscan 7 or an Olympus MVX10 microscope. Quantification of total vascular area, avascular area, and neovascular tufts was performed as previously described^20^.

### Retinal cell death and proliferation assays

Cell death was evaluated using the Click-iT™ Plus TUNEL Assay Kit on paraffin-embedded eye sections. TUNEL-positive cells were quantified using ZEN software. Standardized “Spline regions of interest” were generated, with a 240 μm straight spacer positioned over the optic nerve head. From each end, a 600 μm spline was traced along the curvature of the innermost retinal layer. TUNEL-positive cells were counted along normal vectors projected from this spline. For EC proliferation, dissected retinas were stained with eFluor™ 660-conjugated anti-Ki67 antibody and Alexa Fluor 488-conjugated isolectin GS-IB4. Ki67-positive ECs were quantified using object-based distance measurements in Imaris software (Bitplane, Oxford Instruments) applying a 0 μm distance threshold for colocalization with IB4-labeled ECs. All analyses were blinded and performed across four predefined ROIs per retina. In both assays, the retinal vasculature was visualized using Alexa Fluor 488-conjugated isolectin GS-IB4. All samples were imaged using a Zeiss Axioscan 7 digital slide scanner.

### Primary human retinal microvascular endothelial cell (HRMEC) culture and TXN1 knockdown

HRMECs were obtained from Cell Systems (Cat. #ACBRI 181) and cultured in EBM-2 basal medium (Lonza; Cat. #CC-3156) supplemented with EGM-2 MV Microvascular Endothelial Cell Growth Medium SingleQuots (Lonza; Cat. #CC-4147) on fibronectin (Sigma; Cat. #F1141)-coated dishes. Cells at passages 5-8 were used for all experiments. For TXN1 knockdown, two shRNA constructs targeting the human TXN1 gene (clone ID: TRCN0000064282 and TRCN0000064278) and a non-targeting scrambled control plasmid were obtained from Sigma Aldrich (Mission shRNA Custom DNA). Lentiviral particles were produced in 293T cells^22^ and used to transduce HRMECs. Transduced cells were selected with 1 µg/mL puromycin for three days and used for all downstream assays.

### RT^2^ profiler PCR array

Stably transduced HRMECs were exposed to 85% oxygen for four days. Total RNA was extracted using TRIzol (Invitrogen Cat. #15596026), and cDNA was synthesized with the RT^2^ First Strand Kit (QIAGEN Cat. #330401). Gene expression profiling was performed using the human angiogenesis (PAHS-024Z) and human oxidative stress pathway plus (PAHS-065Y) RT^2^ profiler PCR array from QIAGEN. Quantitative reverse transcription PCR (qRT-PCR) was performed on an ABI QuantStudio 3 system (Applied Biosystems). Selected genes showing significant differential expression in the RT^2^ Profiler PCR array were further validated by additional qRT-PCR. One microgram of total RNA was reverse transcribed using the High-Capacity cDNA Reverse Transcription Kit (Applied Biosystems, Cat. #4368814). Real-time PCR was performed with PowerTrack SYBR Green Master Mix (Applied Biosystems, Cat. #A46109). Relative gene expression was calculated using the comparative ΔCt method and normalized to GAPDH.

## RESULTS

### Generation of EC-specific *Txn1* knockout mice

*Txn1^fl/fl^* mice were generated (Figure 1A) and crossed with *R26-CreER* mice to obtain *Txn1^fl/fl^;R26-CreER* mice. Following tamoxifen administration, *Txn1^fl/fl^;R26-CreER* mice exhibited a significant reduction in TXN1 protein and mRNA levels in lung and brain tissues (Figure 1B,C). To investigate the pathophysiological role of TXN1 during postnatal development of retinal vasculature and retinopathy progression, we generated tamoxifen-inducible, EC-specific *Txn1* knockout mice using *Cdh5-CreERT2* mice (*Txn1^fl/fl^;Cdh5-CreERT2*, hereinafter *Txn1-iECKO*) (Figure 1D). We verified Cre recombination efficiency in retinal ECs under our tamoxifen regimen using *mT/mG;Cdh5-CreERT2* mice, which showed EGFP expression throughout the retinal vasculature at P12 after tamoxifen administration from P5 to P7 (Supplementary Figure S1A).

**Figure 1.**
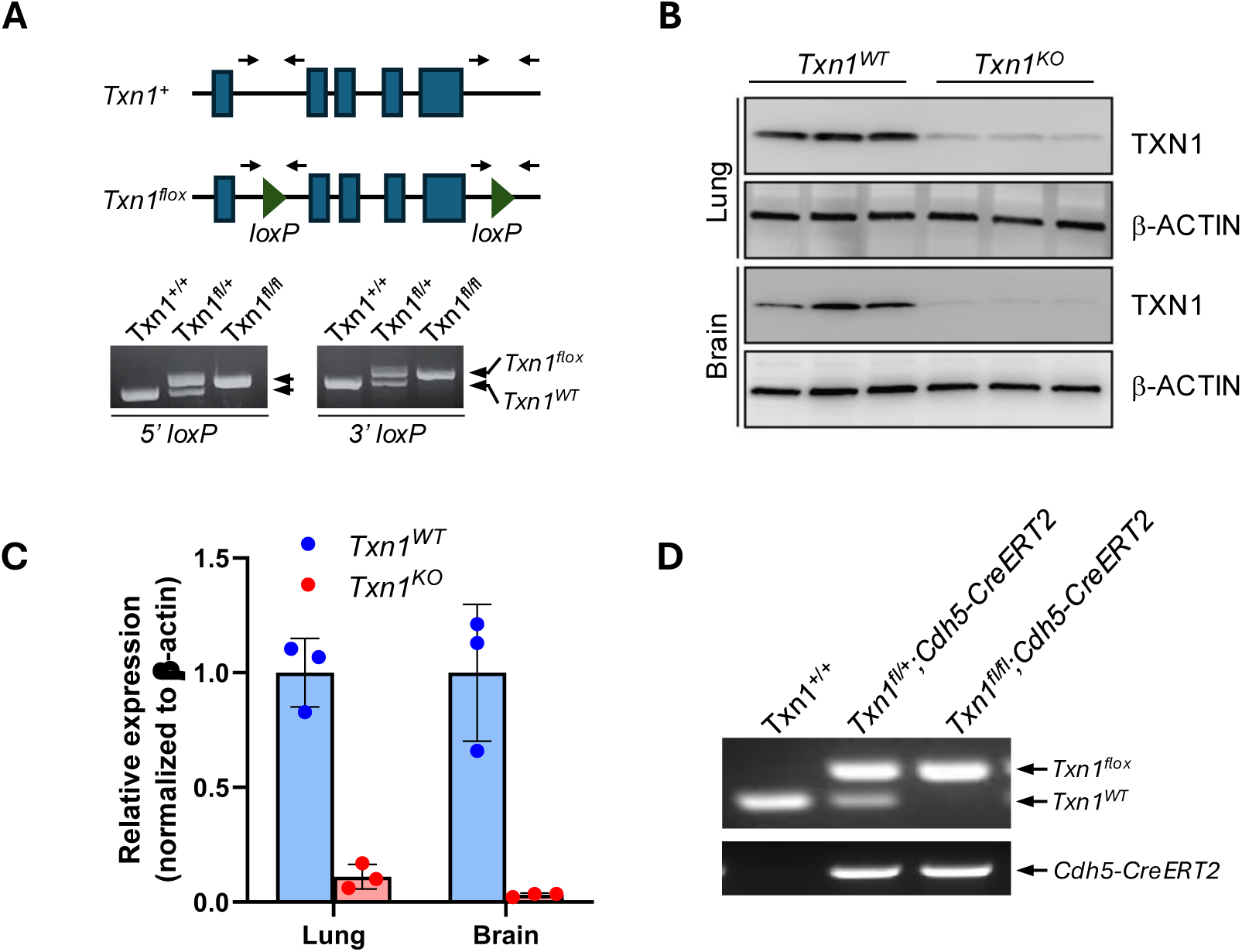
Generation of *Txn1* conditional knockout mice. (A) Schematic representation of wild-type (*Txn1^+^*) and floxed (*Txn1^flox^*) alleles, along with the locations of genotyping primer sets for the 5’ and 3’ loxP sites. (B,C) Validation of TXN1 expression levels in lung and brain tissues from tamoxifen-treated *Txn1^+/+^;R26-CreERT2* (*Txn1^WT^*) and *Txn1^fl/fl^;R26-CreERT2* (*Txn1^KO^*) mice by immunoblotting (B) and RT-qPCR (C). Eight-week-old *Txn1^WT^*and *Txn1^KO^* mice received tamoxifen (100 mg/kg body weight) via intraperitoneal injection for four consecutive days. Lung and brain tissues were collected from three mice per genotype 10 days after the final tamoxifen injection. (D) Genotyping results from *Txn1^+/+^*, *Txn1^fl/+^;Cdh5-CreERT2*, and *Txn1^fl/fl^;Cdh5-CreERT2* mouse tails.

### Deletion of *Txn1* in ECs enhances pathological vaso-obliteration and neovascularization in OIR

The OIR model is a well-established system for recapitulating key pathological features of ROP, including vascular arrest and subsequent neovascularization. Using the mouse OIR model, we investigated the effect of endothelial-specific TXN1 deletion on retinal angiogenesis under hyperoxic conditions. *Txn1^fl/fl^*mice were bred with *Txn1^fl/fl^;Cdh5-CreERT2^+^* mice to obtain *Txn1^fl/fl^* littermates as wild-type controls and *Txn1-iECKO* mice. Both wild-type and *Txn1-iECKO* mice received tamoxifen for three consecutive days (P5-P7) under room-air conditions (Supplementary Figure S1B). We confirmed that body weights did not differ between wild-type and *Txn1-iECKO* mice prior to hyperoxic challenge (Supplementary Figure S2A). In addition to this, evaluation of baseline retinal vascular morphology at P7 revealed that EC-specific deletion of *Txn1* did not cause gross abnormalities in the retinal vasculature (Supplementary Figure S2B). Quantitative assessment of the superficial vascular plexus area further revealed no significant differences between groups (Supplementary Figure S2C), indicating that tamoxifen-induced EC-specific *Txn1* deletion did not affect early retinal vascular development.

We next investigated the specific role of TXN1 in retinal vessel development during the hyperoxia-induced vaso-obliteration phase of OIR. Following tamoxifen treatment from P5 to P7, wild-type and *Txn1-iECKO* mice were exposed to 85% oxygen from P7 to P12, after which retinas were collected at P12 and prepared for vascular assessment. Hyperoxia exposure reduced body weight in the OIR group compared with room-air controls, regardless of genotype; however, no significant differences in body weight were observed between wild-type and *Txn1-iECKO* mice in either condition (Supplementary Figure S3). As expected, both wild-type control and *Txn1-iECKO* retinas exhibited ischemic, avascular regions under hyperoxia at P12 (Figure 2A). Notably, *Txn1-iECKO* retinas showed a significantly reduced vascularized area compared with wild-type controls under the same conditions (Figure 2B), indicating that TXN1-deficient ECs undergo enhanced vascular arrest and regression – hallmark features of the vaso-obliteration phase.

**Figure 2.**
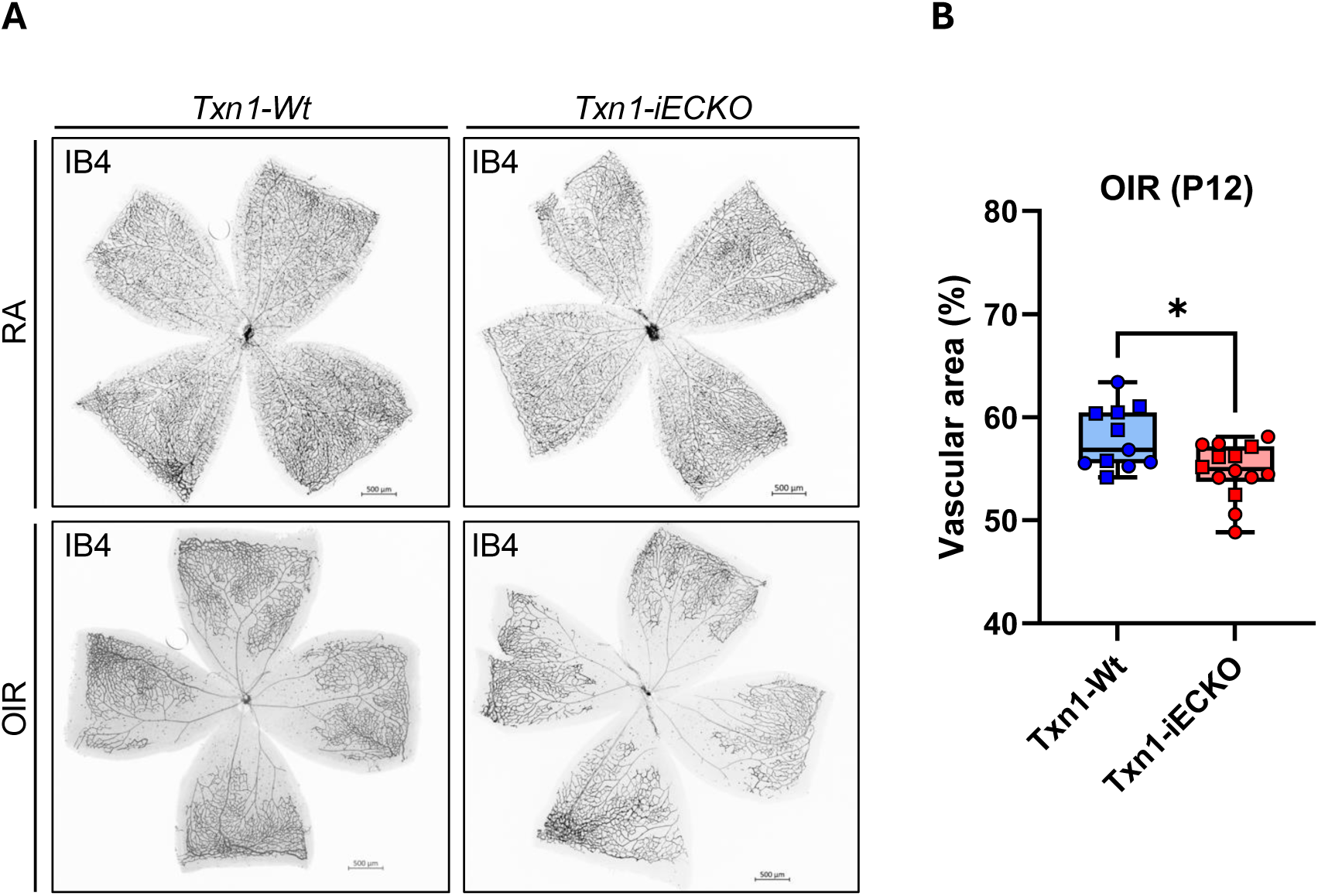
Ablation of Txn1 in ECs results in greater vasoobliteration than in wild-type controls. (A) Representative flat-mount image at P12 showing a reduced vascular area in *Txn1-iECKO* compared to *Txn1-Wt* retinas under hyperoxia (OIR). No gross defects in vessel development were observed in the *Txn1-iECKO* retina under normal room air conditions (RA) compared to the control. Vasculature was visualized by Isolectin B4 staining. Scale bar = 500 μm. (B) Quantification of the percentage of vascular area relative to total retinal area (n=11 for *Txn1-Wt* and n=14 for *Txn1-iECKO*).

Next, we examined neovascularization at P17 following the same hyperoxia treatment protocol. Wild-type and *Txn1-iECKO* mice were returned to room air from P12, and retinas were collected at P17 for vascular assessment. Both wild-type and *Txn1-iECKO* retinas developed extensive neovascular tufts during the neovascularization phase of OIR (Figure 3A). Notably, *Txn1*-deficient retinas exhibited enhanced neovascular tuft formation compared to wild-type controls at this phase (Figure 3B). However, no significant difference was observed in the percentage of vascular area relative to the total retinal area between the two groups, although *Txn1-iECKO* retinas showed a slight reduction (Figure 3C).

**Figure 3.**
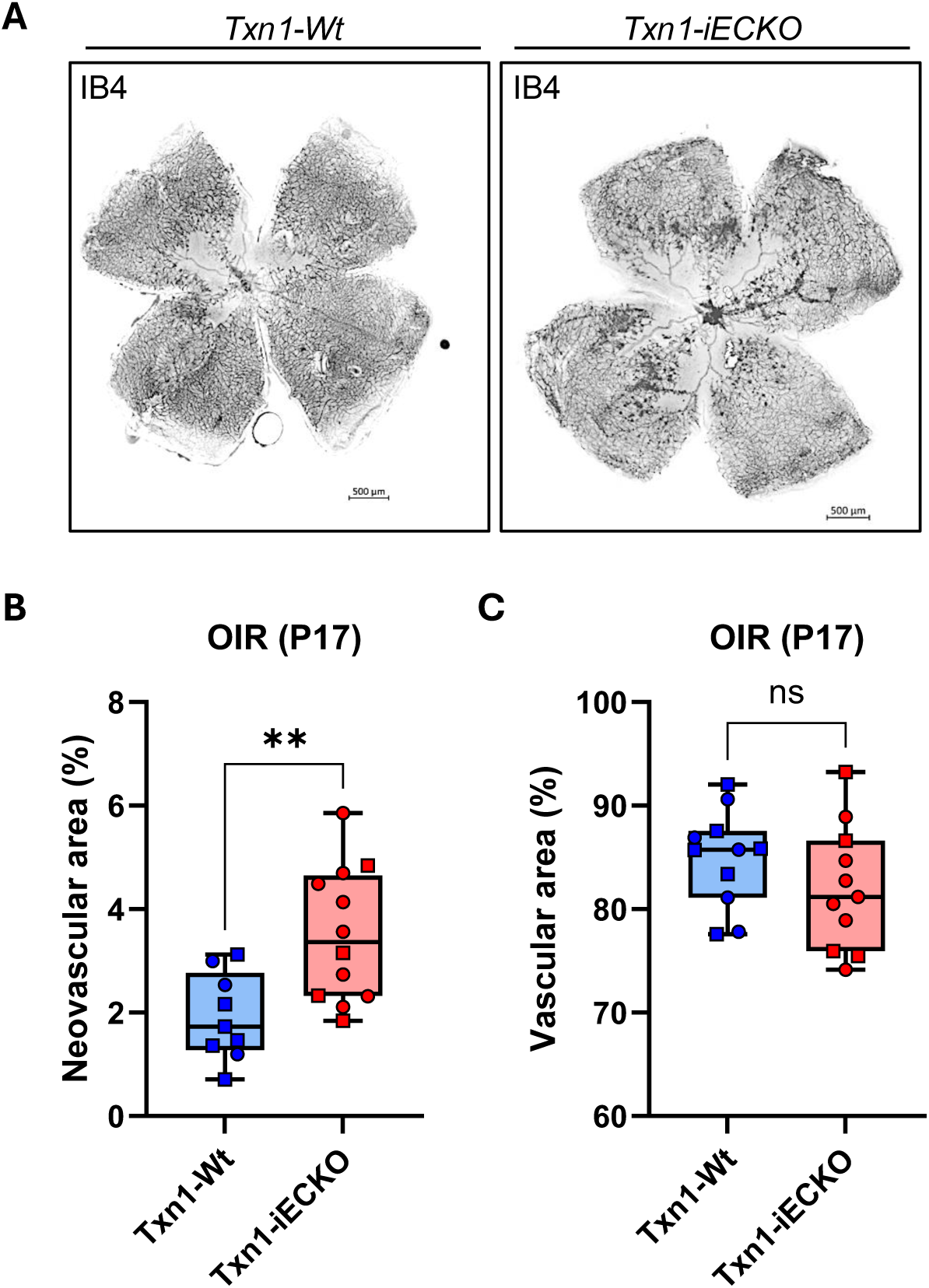
TXN1 protects against pathological neovascularization in the OIR model. (A) Representative images of the neovascularization phase in the OIR model at P17. Retinal vessels were stained with Isolectin B4 antibody. Scale bar = 500 μm. (B, C) Quantification of neovascular tuft formation (percentage of neovascular area) (B) and total vascular area (C) relative to the total retinal area during the neovascularization phase at P17.

### TXN1-deficient ECs show impaired proliferation during vaso-obliteration

Suppression of EC proliferation by hyperoxia is a well-established contributor to the ROP development.^23,24^ During the vaso-obliteration phase, retinal ECs exposed to hyperoxia experience reduced levels of pro-angiogenic factors, including insulin-like growth factor-1 (IGF-1) and vascular endothelial growth factor (VEGF), leading to vascular arrest and vessel regression via EC apoptosis. Given that TXN1 is a known regulator of VEGF production through redox modulation of hypoxia-inducible factor 1α (HIF1α),^25,26^ and that EC-specific deletion of TXN1 resulted in a reduced vascular area at P12 in the OIR model, we hypothesized that this phenotype arises from enhanced vascular regression or arrest, and/or increased apoptotic cell death in *Txn1-iECKO* retinas. To test this, we quantified proliferating ECs by counting Ki67-positive cells colocalized with IB4-labeled vessels. At P7, prior to hyperoxic exposure, no differences in EC proliferation were observed between wild-type and *Txn1-iECKO* retinas under room-air conditions (Supplementary Figure S4A). As expected, hyperoxia exposure reduced EC proliferation compared to room-air conditions, regardless of TXN1 presence (Figure 4A). Under room-air conditions, no differences in EC proliferation were observed between wild-type and *Txn1-iECKO* retinas. However, after five days of hyperoxia exposure from P7 to P12, *Txn1-iECKO* retinas exhibited a significant reduction in Ki67-positive proliferating ECs compared with wild-type controls (Figure 4A).

**Figure 4.**
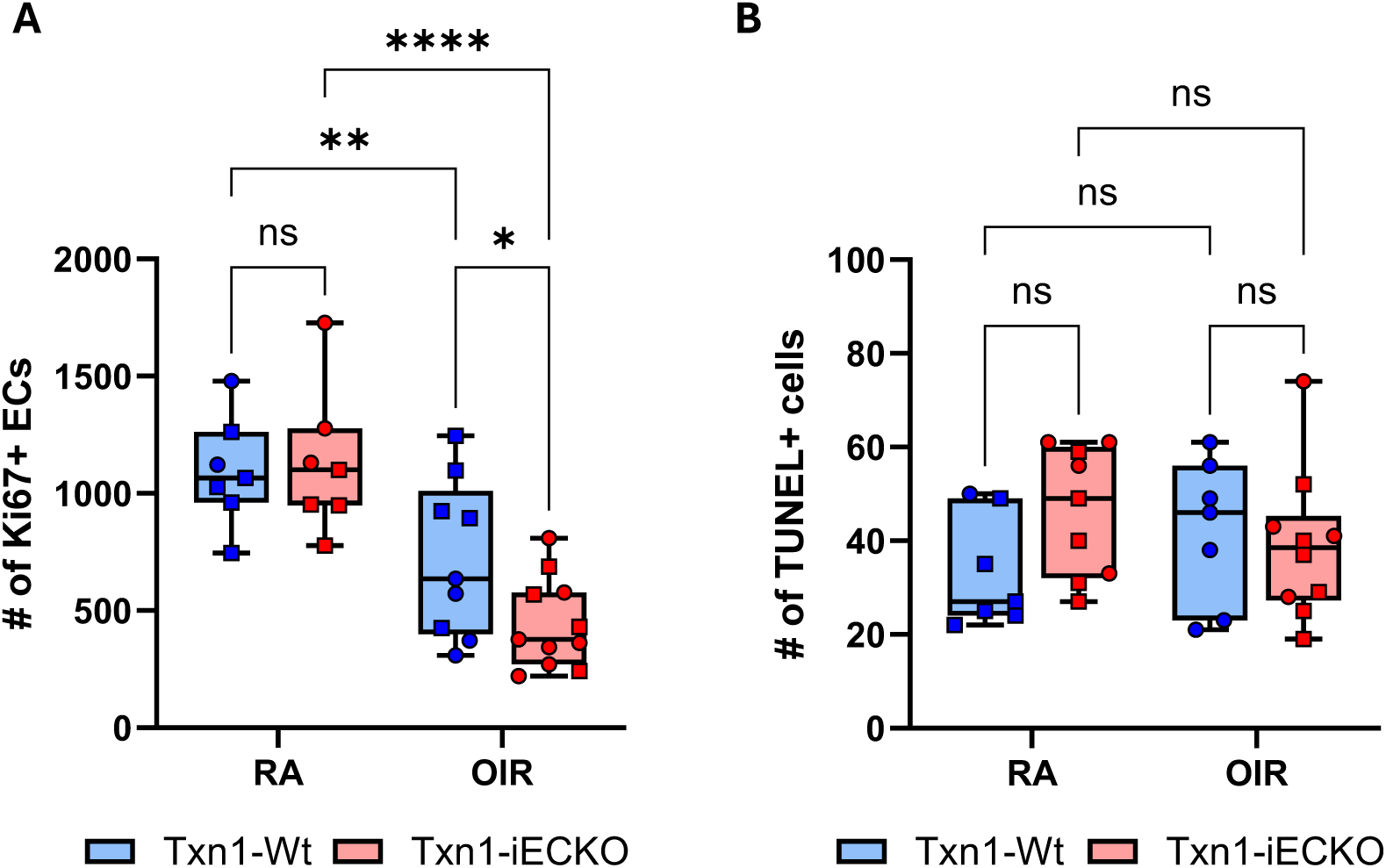
Loss of TXN1 significantly affects retinal EC proliferation but not survival. (A) Quantification of Ki67-positive ECs in flat-mounted retinas from *Txn1-Wt* and *Txn1-iECKO* mice at P12 under RA and OIR conditions (*Txn1-Wt*: n=7, *Txn1-iECKO*: n=7 in the RA group; *Txn1-Wt*: n=9, *Txn1-iECKO*: n=11 in the OIR group). (B) Quantification of TUNEL-positive cells in retinal sections from *Txn1-Wt* and *Txn1-iECKO* mice at P12 under RA and OIR conditions (*Txn1-Wt*: n=7, *Txn1-iECKO*: n=9 in the RA group; *Txn1-Wt*: n=7, *Txn1-iECKO*: n=10 in the OIR group).

Given the role of TXN1 in redox homeostasis and limiting oxidative stress-induced apoptosis^27,28^, we hypothesized that *Txn1-iECKO* retinas would exhibit increased cell death. Using the same OIR protocol, retinas from wild-type and *Txn1-iECKO* mice were harvested at P7 and P12 and analyzed for apoptosis by TUNEL staining. No differences in apoptotic cell death were observed between genotypes at P7 room-air conditions (Supplementary Figure S4B). Furthermore, at P12, exposure to hyperoxia did not increase retinal apoptosis compared with room-air controls, and EC-specific deletion of TXN1 had no detectable effect (Figure 4B). Collectively, these results indicate that TXN1 deletion exacerbates hyperoxia-induced vascular regression by impairing EC proliferation, rather than by increasing overall retinal cell death at P12, potentially due to compensation by other antioxidant systems in the retina.

### TXN1 knockdown activates NRF2 signaling in HRMECs under hyperoxia

Given the established protective role of TXN1 against reactive oxygen species,^27,28^ we further investigated TXN1-dependent transcriptional changes in ECs under normoxic and hyperoxic conditions. HRMECs were transduced with two independent shRNA-expressing lentiviruses targeting TXN1. After three days of puromycin selection, we confirmed efficient TXN1 knockdown in both shTXN1-infected HRMECs by immunoblotting and quantitative RT-PCR (Figure 5A,B). No significant difference in cell proliferation was observed between shTXN1-HRMECs and scrambled shRNA controls under normoxic conditions (data not shown).

**Figure 5.**
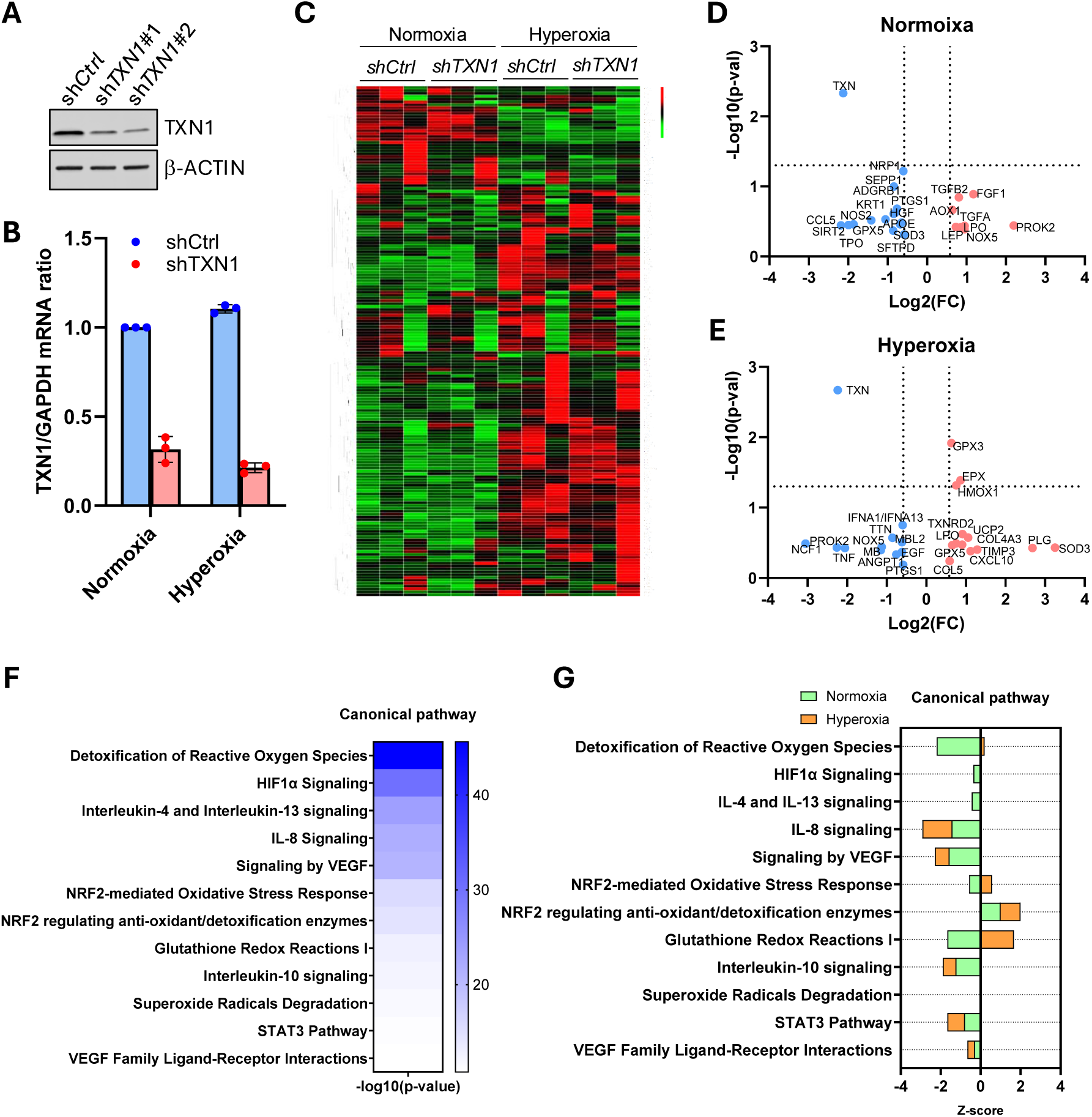
TXN1-mediated alterations in gene expression and the enrichment of associated pathways in HRMECs. (A, B) Immunoblotting (A) and quantitative RT-PCR (B) analysis confirming the efficiency of shRNA-mediated TXN1 knockdown in HRMECs. (C) A one-way hierarchical heatmap was created by combining the 2^(-dCt) fold change values from the Angiogenesis and Oxidative Stress Pathway Plus array panels using R version 4.4.1, along with the heatmap package. Fold change and p-values were obtained from the RT2 Profiler PCR Array Data Analysis Spreadsheet 1904 provided by QIAGEN. The expression values were Z-score normalized within a range of -3 to 3 to indicate the degree of down-regulation or up-regulation. (D, E) Volcano plots were generated from the normoxia (D) and hyperoxia (E) -treated groups, using a 1.5-fold change cutoff indicated by the vertical dotted lines. A p-value of less than 0.05 is represented by the horizontal dotted line based on data from all genes in (C). (F, G) Ingenuity Pathway Analysis (IPA) shows significantly enriched canonical pathways (F) and predicts downstream effects of activation (positive Z-scores) or repression (negative Z-scores) (G) in the pathways of sh*TXN1*-HRMECs.

To examine the downstream effects of TXN1 under hyperoxia, we exposed shTXN1- and control HRMECs to 85% or 21% oxygen for four days to simulate hyperoxic and normoxic conditions, respectively. Differential gene expression was analyzed using RT^2^ Profiler PCR arrays targeting human angiogenesis and oxidative stress pathways. Heatmap analysis revealed distinct transcriptional changes in both angiogenesis and oxidative stress pathways between normoxic and hyperoxic conditions (Figure 5C). Although most genes did not reach statistical significance, except for TXN1 (p=0.005), TXN1 knockdown appeared to modulate key genes in angiogenesis and oxidative stress pathways. Under normoxia, TXN1-deficient HRMECs showed upregulation of pro-angiogenic and redox-related genes, including PROK2, FGF1, TGFA, TGFB2, and NOX5, and downregulation of NRP1, ADGRB1, PTGS1, NOS2, GPX5, and SOD3 (Figure 5D). Under hyperoxia, TXN1 knockdown induced a distinct profile, with upregulation of antioxidant genes GPX3, EPX, HMOX1, TXNRD2, SOD3, and GPX5, and downregulation of NCF1, PROK2, NOX5, and ANGPT1 (Figure 5E). These findings indicate that loss of TXN1 reprograms endothelial gene expression in an oxygen-dependent manner.

To gain functional insight into the observed transcriptional changes, we performed Ingenuity Pathway Analysis (IPA) on the differentially expressed genes under normoxic and hyperoxic conditions. We identified significant enrichment of canonical pathways, including HIF1α signaling, VEGF signaling, and the nuclear factor erythroid 2-related factor 2 (NFE2L2/NRF2)-mediated oxidative stress response pathways, in shTXN1-HRMECs compared with control cells under both normoxic and hyperoxic conditions (Figure 5F). Notably, TXN1 knockdown modulated several oxidative stress-related pathways in a context-dependent manner. Under normoxia, HIF1α signaling, ROS detoxification, and IL-4/IL-13 signaling were suppressed, whereas these pathways were not significantly altered by TXN1 loss under hyperoxia (Figure 5G). In contrast, IL-8, IL-10, STAT3, and VEGF signaling were consistently inhibited by TXN1 knockdown under both normoxic and hyperoxic conditions. Moreover, the NRF2-mediated oxidative stress response and glutathione redox pathways were activated under hyperoxia but suppressed under normoxia following TXN1 knockdown, suggesting that TXN1 functions as an oxygen-dependent molecular switch in ECs (Figure 5G).

Given the established role of NRF2 in regulating thioredoxin reductase 1 (TXNRD1)^29,30^ and TXNIP,^16,31,32^ as well as its crucial protective function against oxidative stress, we validated the activated NRF2 pathway following TXN1 depletion using quantitative RT-PCR in shTXN1- and control HRMECs. NRF2 expression was significantly upregulated by hyperoxia, independent of TXN1 knockdown (Figure 6). In contrast, expression of the canonical NRF2 target genes NQO1, HMOX1, and TXNRD1 was markedly increased in TXN1-deficient cells under both normoxic and hyperoxic conditions, with a more pronounced increase under hyperoxia (Figure 6). These findings indicate that TXN1 knockdown enhances NRF2-mediated antioxidant pathways in ECs, particularly under oxidative stress.

**Figure 6.**
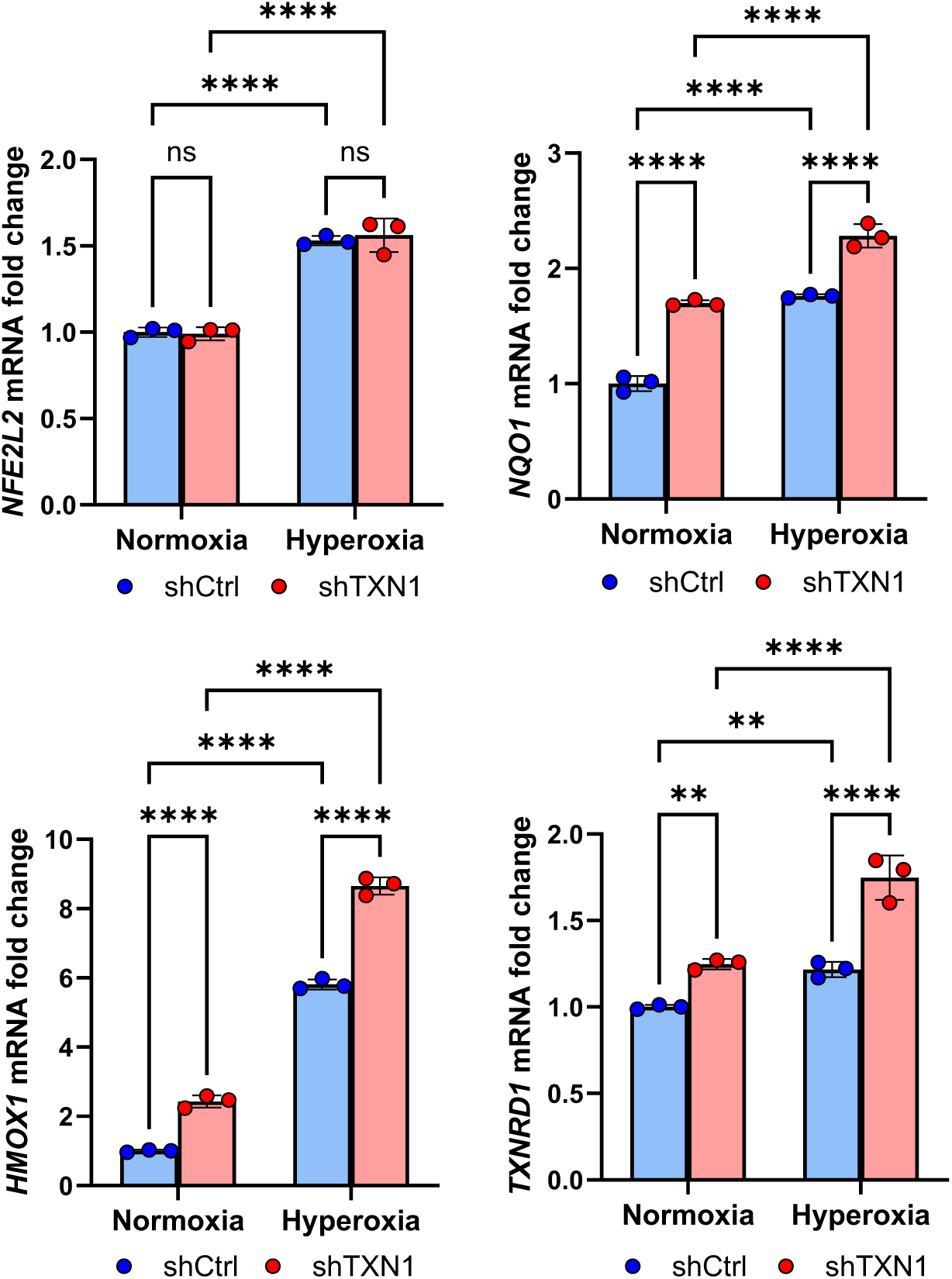
Quantitative RT-PCR (qRT-PCR) analysis of sh*TXN1* or control shRNA-HRMECs was conducted. Cells were cultured under hyperoxic (85% O₂) or normoxic (21% O₂) conditions for four days and then analyzed by qRT-PCR across two independent experiments. One of the representative data is presented as the mean of triplicate PCR reactions. Statistical significance was determined using two-way ANOVA: ****p < 0.0001, ***p = 0.0001, **p < 0.01.

## DISCUSSION

In this study, we identify TXN1 as an essential regulator of vascular development, particularly in the context of ischemic retinopathy. Using the OIR mouse model, we demonstrate that endothelial cell-specific deletion of TXN1 enhances vascular regression during the hyperoxic vaso-obliteration phase and significantly exacerbates pathological angiogenesis, particularly neovascular tuft formation, during the subsequent hypoxic neovascularization phase. TXN1-deficient retinas showed a significant reduction in endothelial proliferation without corresponding changes in TUNEL-positive retinal cells, indicating that impaired vascular growth, rather than increased apoptosis, drives the vascular abnormalities observed under hyperoxic conditions. Transcriptomic profiling of TXN1-deficient HRMECs exposed to hyperoxia further revealed increased expression of NRF2-dependent genes, underscoring a central role for the thioredoxin system in modulating the endothelial oxidative stress response.

A previous study by Ozawa et al. demonstrated that TXN1 overexpression protects the retinal vasculature from hyperoxia-induced injury in the OIR model.^18^ Transgenic mice expressing human TXN1 (TXN-Tg) exhibited an expanded vascularized area during the vaso-obliteration phase and reduced neovascular tuft formation during the neovascularization phase. Our EC–specific *Txn1* knockout model further supports TXN1’s protective role, particularly in regulating vascular development under hyperoxic stress. In *Txn1-iECKO* retinas, we observed more pronounced vascular regression during the vaso-obliteration phase and increased pathological neovascularization during the neovascular phase. These phenotypes parallel the findings of Ozawa et al., but our results establish that these effects arise specifically from endothelial loss of TXN1.

Several mechanisms could explain the vascular phenotypes seen in the *Txn1-iECKO* retinas. It is possible that transient or early endothelial cell death occurred before P12 in the OIR model, leading to a missed detection window in our analysis. Future studies using earlier time points or early apoptotic markers, such as cleaved caspase-3, could better capture the dynamics of endothelial cell death.

Alternatively, the increased vascular regression in *Txn1-iECKO* retinas may reflect impaired angiogenic capacity rather than enhanced apoptosis. TXN1 could promote EC sprouting and migration, particularly under oxidative stress, potentially through pathways involving VEGFR2, DLL4,^33^ or CXCR4.^34^ From another perspective, TXN1 deficiency may delay, rather than inhibit, angiogenesis. Transient reductions in key proangiogenic factors such as VEGF-A or TGF-β1 could account for the delayed vascularization observed at P12, with potential recovery at later neovascular stages. Moreover, TXN1 may regulate vascular remodeling or regression^35,36^ through apoptosis-independent mechanisms, such as endothelial cell retraction,^37^ basement membrane remodeling,^38,39^ and microglia-mediated pruning,^40–42^ either directly or indirectly, thereby reducing vascular density without altering proliferation or cell death. Together, these observations indicate that TXN1 contributes to retinal vascular development and stability during OIR not only by maintaining redox homeostasis via HIF-1α-mediated regulation of VEGF,^43^ but also by influencing angiogenesis and vascular remodeling.

Our findings further suggest a dynamic interplay between the TXN system and NRF2 signaling during ROP progression. We observed that TXN1 knockdown in HRMECs triggers NRF2 activation, as evidenced by upregulation of canonical target genes such as NQO1, HMOX1, and TXNRD1, with this response being more pronounced under hyperoxic conditions, where oxidative stress is elevated. NRF2 functions as a central regulator of the cellular antioxidant defense and maintain redox homeostasis.^30,44–46^ Given TXN1’s role in maintaining antioxidant systems, disruption of TXN system leads to compensatory upregulation of other antioxidant pathways, such as glutathione peroxidase (GPX), peroxiredoxin (PRDX), and sulfiredoxin (SRX), via NRF2 activation^47^ to restore redox balance.

Beyond NRF2 activation, TXN1 deficiency reshapes broader oxidative stress–related signaling networks in a context-dependent manner. Under normoxia, depletion of TXN1 suppresses protective pathways including HIF1α signaling, ROS detoxification, and IL-4/IL-13 signaling, while hyperoxia leaves these pathways unchanged. In contrast, cytokine and growth factor pathways such as IL-8, IL-10, STAT3, and VEGF signaling are consistently inhibited regardless of oxygen tension.

Taken together, our findings suggest that TXN1 functions as a molecular switch, regulating endothelial antioxidant defenses in an oxygen-dependent manner while contributing to cellular susceptibility to oxidative stress in ROP. Furthermore, our results confirm that a robust and redundant antioxidant network regulated by NRF2 can compensate for the loss of TXN1, maintaining redox balance and supporting EC resilience.

## Acknowledgements

The authors would like to thank Mark Dittmar (Dean McGee Eye Institute, OK) and Ben Fowler (Oklahoma Medical Research Foundation, OK) for their invaluable support with animal experiments and imaging. During the preparation of this work, the author(s) used Jenni AI (2025 Jenni, Inc.) in order to improve the clarity and quality of written communication with grammar checking and sentence restructuring. After using this tool/service, the author(s) reviewed and edited the content as needed and take full responsibility for the content of the publication.

## Commercial Relationships Disclosure

None

## CONFIRMATION OF THE AUTHOR’S PUBLICATION AND LICENSING RIGHTS

Science Suite Inc. dba BioRender (“BioRender”) has granted Faizah Bhatti permission to use this Completed Graphic in accordance with BioRender’s Terms of Service and Academic License Terms (“License Terms”). Bhatti, F. (2025) https://BioRender.com/8a8edya

---

**Supplementary Figure S1**. Optimization of tamoxifen inducible OIR model. (A) *mT/mG;Cdh5-CreERT2* reporter mice were generated by crossing *mT/mG mice* (The Jackson Laboratory) with *Cdh5-CreERT2* mice. 50 mg of tamoxifen was administered via oral gavage for three consecutive days from P5 to P7, and the retina was harvested and flat-mounted at P12. These reporter mice were utilized to optimize tamoxifen administration and dosing, as well as to further confirm Cre recombinase activity in the retinal vessels of experimental *Txn1^fl/fl^;Cdh5-CreERT2* mice. (B) Schematic of the oxygen-induced retinopathy mouse model used in this study. Tamoxifen was administered on postnatal days P5, P6, and P7 in wild-type control (*Txn1^fl/fl^*) and *Txn1-iECKO* mice, followed by five consecutive days of hyperoxic treatment (75% oxygen). Retinal vessel development was assessed at P12 and P17 using retinal flat-mount immunohistochemistry staining. Created in BioRender. Bhatti, F. (2025)

**Supplementary Figure S2**. Tamoxifen-induced depletion of *Txn1* in ECs does not significantly affect early retinal vessel development under normoxic room-air conditions. (A) Body weight measurements at P7 show no significant difference between *Txn1-iECKO* (n=7) and wild-type control (n=9) mice following *Txn1* ablation. (B) Representative images of retinal flat-mounts from wild-type control (*Txn1-Wt*) and *Txn1-iECKO* at P7. Retinal vessels were visualized using anti-CD31 antibody staining. Scale bar = 500 μm. (C) The percentage of vascular area relative to total retinal area shows no significant difference between the groups at P7 (n=8 for *Txn1-Wt* control and n=6 for *Txn1-iECKO*).

**Supplementary Figure S3.** Body weight measurements at P12 show no significant difference between wild-type control and *Txn1-iECKO* (*Txn1-Wt*: n=12, *Txn1-iECKO*: n=17 in the RA group; *Txn1-Wt*: n=18, *Txn1-iECKO*: n=15 in the OIR group).

**Supplementary Figure S4.** (A) Quantification of Ki67-positive ECs in flat-mounted retinas from *Txn1-Wt* and *Txn1-iECKO* mice at P7 under room-air conditions (*Txn1-Wt*: n=5, *Txn1-iECKO*: n=3). (B) Quantification of TUNEL-positive cells in retinal sections from *Txn1-Wt* and *Txn1-iECKO* mice at P7 under the same conditions (*Txn1-Wt*: n=12, *Txn1-iECKO*: n=10).

