## Supplemental Data for "Endothelial Cell Specific Thioredoxin 1-Dependent Signaling in Oxygen-Induced Retinopathy"

### Supplementary Figure S1

A

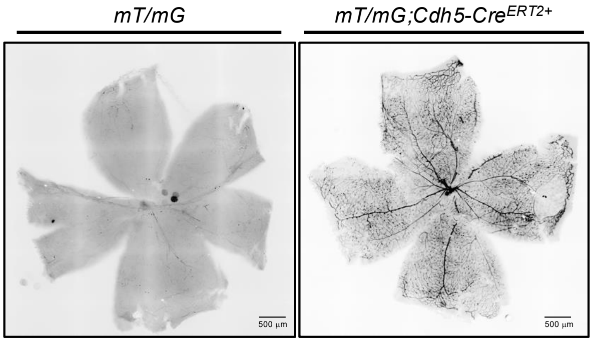


B

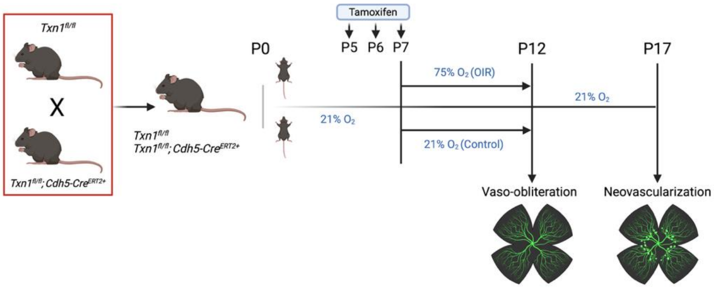

### Supplementary Figure S2

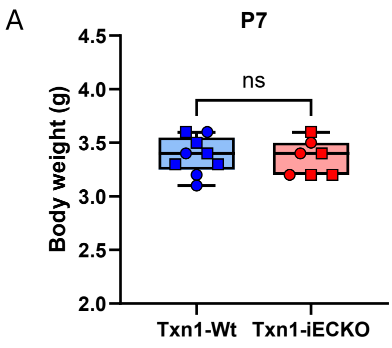


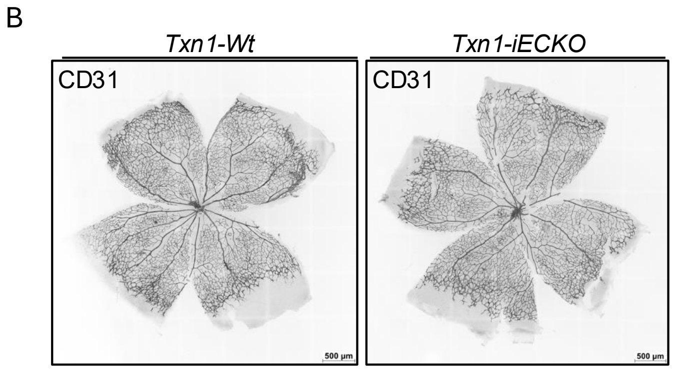


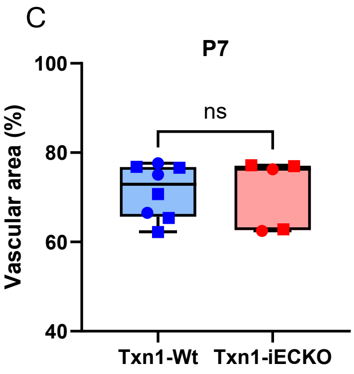

### Supplementary Figure S3

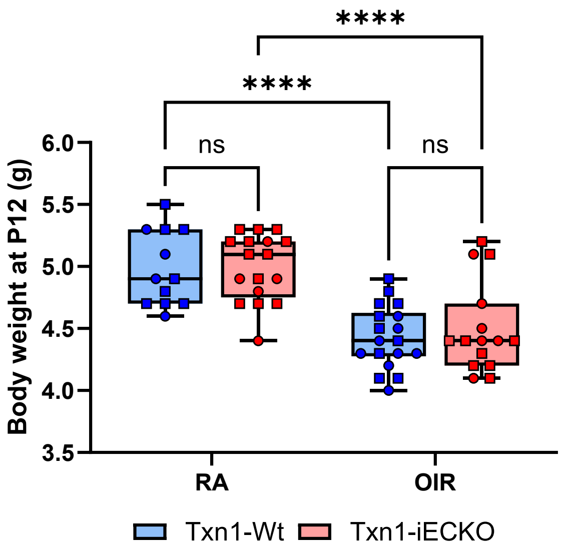

### Supplementary Figure S4

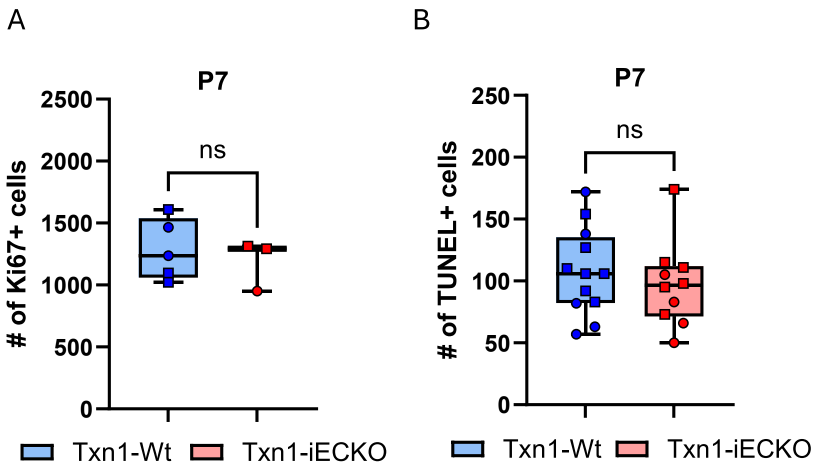
